# Influence of distractors on spatial working memory and neural activity in marmoset prefrontal cortex

**DOI:** 10.64898/2026.05.01.722299

**Authors:** Raymond Ka Wong, Janahan Selvanayagam, Kevin D. Johnston, Alessandro Zanini, Margaret Sadie Loewith, Stefan Everling

## Abstract

The prefrontal cortex (PFC) plays a critical role in maintaining working memory (WM) representations while filtering irrelevant distractors. In macaques, PFC neurons exhibit persistent delay period activity that is robust to distractor interference. The common marmoset has emerged recently as a complementary primate model for investigating the neural basis of cognitive processes including WM, in part because the relatively lissencephalic cortex of this species enables laminar recordings which could provide substantial insight into the microcircuit basis of these functions. It remains unknown however, whether marmoset WM performance is robust to distractors presented during delay periods of WM tasks, and how such distractor filtering may be implemented in PFC circuits. Here, we addressed this gap by conducting wireless recordings of PFC in freely moving marmosets performing a touchscreen-based delayed-match-to-location (DML) task in which a salient visual distractor was presented during the delay period on a subset of trials. Marmosets maintained WM performance on distractor trials, showing a decrease in accuracy of only 5%. Consistent with prior observations in both the macaque and marmoset models, we found that many PFC neurons exhibited activity related to the stimulus sample, during the delay period, and around the time of the behavioural response. In a subset of neurons, we observed distractor-mediated modulations of persistent delay period activity which were associated with a greater incidence of performance errors on the DML task. These findings reveal that marmoset WM is robust to distractor interference, and that the PFC mechanisms instantiating WM and distractor filtering are conserved in this primate species. Taken together, they support the common marmoset as a complementary model for investigating the contribution of PFC circuits to mnemonic and attentional processes.

## Introduction

Working memory (WM), the ability to temporarily maintain and manipulate information in the absence of sensory input, is a cognitive process critical to the orchestration of goal-directed behavior and higher-order cognition (D’Esposito and Postle 2015). A core aspect of WM function is resistance to distractor interference. In order to function effectively in complex environments, WM representations must be robust to the incoming barrage of incoming sensory information irrelevant to the task at hand (Lorenc et al. 2021). Elucidating the neural mechanisms by which the brain filters such distractors is essential to understanding how the brain implements flexible, interference-resistant WM.

Persistent delay-period activity in the prefrontal cortex (PFC) has long been identified as a critical neural correlate of working memory (Fuster and Alexander 1971; Goldman-Rakic 1995). A series of now classic investigations combining performance of an oculomotor delayed response (ODR) task with single-neuron recordings in rhesus macaques established that dorsolateral PFC (dlPFC) neurons exhibit sustained, spatially selective activity during a delay period intervening between the brief presentation of a visual stimulus and the subsequent response to the remembered stimulus location (Funahashi et al. 1989; Chafee and Goldman-Rakic 1998; Chafee and Goldman-Rakic 2000). Such activity was diminished or truncated on trials on which monkeys made performance errors (Funahashi et al. 1989), linking the discharge rates and dynamics of dlPFC persistent activity with WM performance. Subsequent studies built on these seminal findings to investigate how PFC persistent activity and WM performance are impacted by the presentation of interfering stimuli. When salient distractors are flashed during the delay period of the ODR task, dlPFC neurons exhibit transient disruptions of persistent activity, the magnitude of which correlates with performance errors (Suzuki and Gottlieb 2013; Qi et al. 2021). Reversible inactivation of dlPFC produces substantial decrements in performance on distractor trials, demonstrating a causal role of dlPFC in distractor suppression (Suzuki and Gottlieb 2013). Taken together, these findings suggest that the dlPFC plays a critical role in both the retention of mnemonic information and the suppression of distractor interference.

The common marmoset has recently emerged as a complementary primate model for investigating the neural basis of cognitive processes (Mitchell and Leopold 2015; Mitchell et al. 2016) and may hold particular promise for electrophysiological investigations of the microcircuit mechanisms underlying WM performance and distractor filtering. Current computational models of the instantiation of these processes in PFC circuits make layer and neuron-type specific predictions which could be informed by laminar recordings(Wang et al. 2004; Joyce et al. 2025; see for review Wang 2021). The smooth marmoset PFC is readily accessible to laminar recordings which are practically limited for some regions of macaque PFC which lie within the principal sulcus. Though some species differences exist (Magrou et al. 2026), marmosets and macaques additionally share a suite of homologous PFC subregions which can support our understanding of primate PFC function more broadly(Burman et al. 2006; Reser et al. 2013). Here, the small size and smooth cortex of the marmoset are additionally advantageous, as implanted electrode arrays can cover multiple PFC subregions, allowing simultaneous sampling of neuronal activity during task performance (Wong et al., 2024).

Marmosets have demonstrated spatial WM capabilities across multiple paradigms (Miles 1957; MacDonald et al. 1994; Tsujimoto and Sawaguchi 2002; Spinelli et al. 2004; Spinelli et al. 2006; Nakako et al. 2013; Yamazaki et al. 2016; Sadoun et al. 2019; Glavis-Bloom et al. 2022; Glavis-Bloom et al. 2023; Vanderlip et al. 2025 Mar; Vanderlip et al. 2026 Jan 1), and recent work using touchscreen-based delayed match-to-location (DML) tasks has confirmed the presence of persistent delay-period activity in marmoset PFC (Wong et al. 2023; Wong et al. 2024). It remains unknown however, whether marmosets are able to effectively perform spatial WM tasks in the face of distraction, and thus whether they are a viable model for neurophysiological investigations of distractor interference. Indeed, behavioural evidence of high distractibility, together with established neuroanatomical constraints on resistance to distraction confirmed via modelling, suggest that the marmoset model may be of limited utility for studies of the neural basis of this aspect of WM performance (Joyce et al. 2025; Magrou et al. 2026). To address this, we trained marmosets to perform a touchscreen-based DML task in which a salient distractor was flashed at variable locations during the delay period on a subset of trials, and subsequently carried out simultaneous wireless recordings of neurons across a broad area of the marmoset PFC using chronically implanted 96-channel Utah arrays during performance of this task. We show that marmosets are able to maintain spatial working memory in the face of distraction, with presentation of distractors inducing a moderate decrement in performance on the DML task. Consistent with previous work (Wong et al. 2023) we observed spatially tuned persistent activity during the delay period of the DML task, and extend that finding to show that, similar to previous work using the ODR task in rhesus macaques (Suzuki and Gottlieb 2013), distractor-related disruptions of persistent activity correlated with reduced task performance. Our findings establish the viability of the marmoset model for investigating the neural basis of distractor filtering during WM, and suggest conserved mechanisms of interference control between New and Old World primates.

## Methods

### Subjects

Data were collected from two adult female common marmosets (*Callithrix jacchus*; Marmoset H, 37-42 months; Marmoset D, 37-39 months). All experimental procedures conducted were in accordance with the Canadian Council of Animal Care policy on the care and use of laboratory animals and a protocol approved by the Animal Care Committee of the University of Western Ontario Council on Animal Care. The animals were under the close supervision of university veterinarians.

### Training protocol/Task

Marmosets performed a modified version of the DML task reported in Wong and colleagues (2023) on an in-house developed touchscreen testing box attached to the home cage (Figure 1). Animals were trained through a sequence of phases using successive approximations to establish performance on the task. Each trial began with the presentation of a sample stimulus (filled blue or pink circle, 3 cm diameter) on a gray background at one of the four corner locations of the touchscreen display for 2.5s, followed by a delay period. Choice stimuli then appeared at all four locations, and the animal was required to touch the location matching the sample stimulus to obtain a liquid reward (0.075–0.1 ml of a 50/50 mixture of 1:1 acacia gum powder and water with liquid marshmallow). A detailed explanation of the training phases is reported in Wong et al. (2023). Once marmosets were fully trained on the task with a 4s delay, a task initiation phase was added at the beginning of each trial (see Figure 1). This element was added to the task to ensure active engagement of the animals at the beginning of each trial, analogous to the fixation requirement typically implemented in the ODR task. In this phase, a large visual stimulus (6 cm diameter circle) differing in colour from the sample stimuli, was presented at the center of the touchscreen. When the animal touched this central stimulus, the DML task commenced after a variable delay of 100-500ms. If the animals task performance was deleteriously affected by the addition of this self-initiation phase, we reduced the delay period duration from 4s to 1s to reduce overall task difficulty. Once performance recovered, we incrementally increased the delay back to 4s. Animals were able to complete the initial training protocol in 6-8 weeks, with an additional 1-2 weeks to begin self-initiating the task.

**Figure 1.**
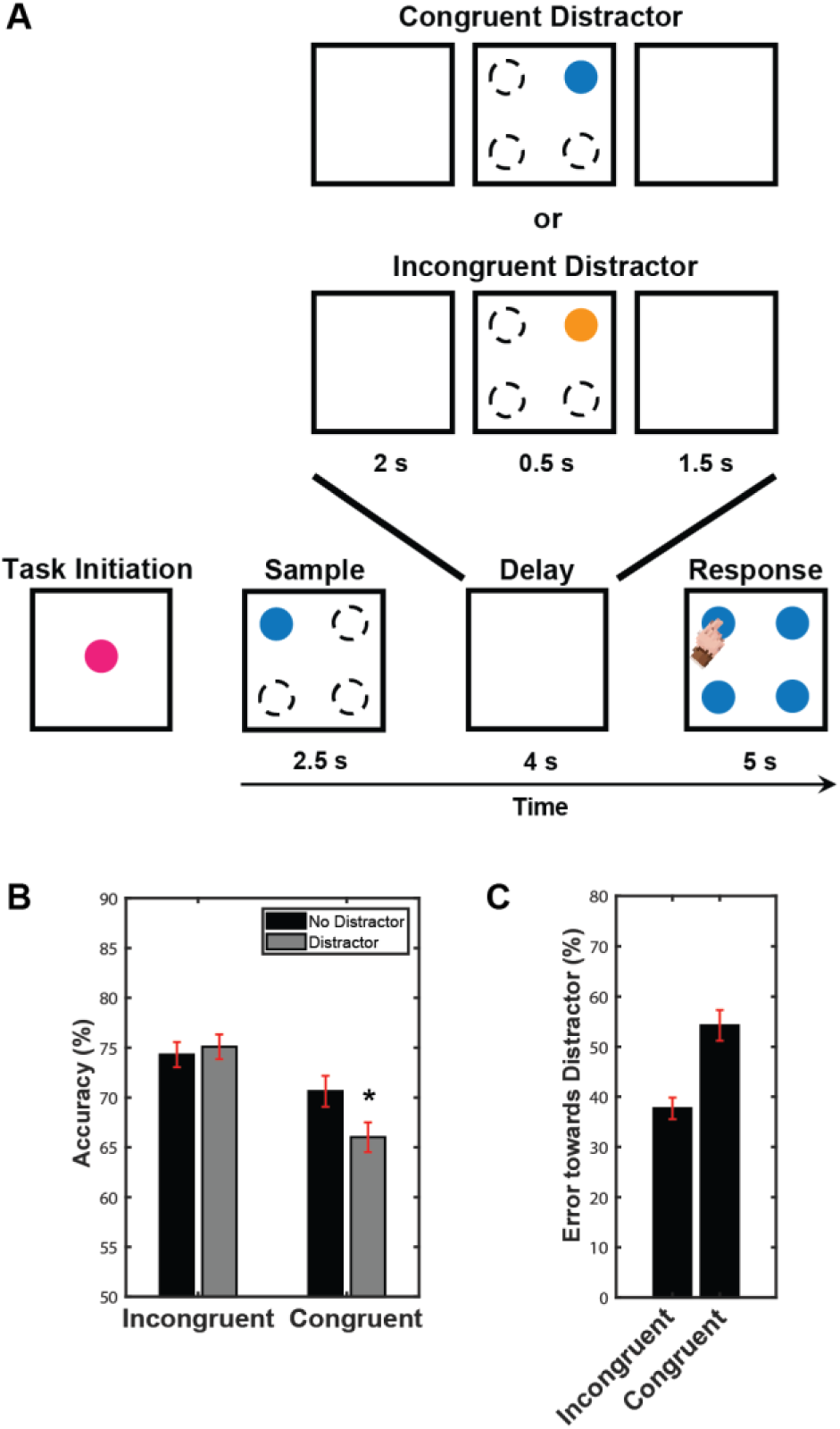
Task and behavioural performance. **A)** DML task. After self-initiation of the task by pressing a central stimulus, successive panels indicate initial appearance of the sample stimulus, delay period and response phases. The sample was randomly presented at 1 of 4 possible locations. On distractor trials (30-50% of trials within each session), a distractor was briefly presented during the delay period at 1 of 3 locations different from that at which the sample stimulus was presented. Within a given session, only incongruent or congruent colour distractors were presented (mutually exclusive). Lower panels, task performance **(B)** Task accuracy for distractor and no-distractor trials for colour-incongruent and colour-congruent distractors. **C)** Proportion of errors directed toward the distractor location for colour-incongruent and colour-congruent distractor types.

During neural recordings, animals performed a version of the self-initiated DML task described above that included a subset of trials in which a single distractor stimulus was presented during the delay period, 2s after sample offset. The distractor stimuli consisted of 6 cm circles and were classified as one of two types. “Congruent” distractors were the same colour as the sample stimulus, whereas “incongruent” distractors differed in colour. These stimuli were presented for 500 ms at one of the three locations not corresponding to the sample stimulus. Distractor trials were pseudorandomly interleaved with no-distractor trials, and occurred on 30-50% of trials within a given session. Distractor type was varied across sessions but remained constant within a session.

### Array surgery

Animals underwent an aseptic surgical procedure under general anesthesia in which 96-channel electrode arrays (4 mm x 4 mm; 1.0 mm electrode length; 400 µm pitch; iridium oxide tips) (Blackrock Neurotech, Salt Lake City, US) were implanted in the left PFC (see Selvanayagam et al. 2019 for details). Briefly, during this surgery, a microdrill was used to perform a ∼5 mm craniotomy which was enlarged as necessary using a rongeur. The dura was carefully opened and the array was manually inserted into the lateral PFC. Wires and connectors were fixed to the skull using dental resin cement (All-Bond Universal and Duo-Link, Bisco Dental Products). Once implanted, the array site was covered with a thin layer of silicone adhesive (Kwik Sil; World Precision Instruments) and then covered with dental cement. A screw hole was drilled into the skull, into which we inserted a stainless-steel ground screw. The ground wire of the array was then tightly wound around the base of the screw to ensure a stable electrical connection. A combination recording chamber/head holder (Johnston et al. 2018) was placed around the array and connectors and fixed in place using further layers of dental resin cement. Finally, a removable protective cap was placed on the chamber to protect the 3×32 channel Omnetics connector.

### Neural recordings

After recovery from array implantation, we verified that electrode contacts were within the cortex by monitoring extracellular neural activity using the SpikeGadgets’ data acquisition system (SpikeGadgets, San Francisco, US). Upon observing single- or multiunit activity at multiple sites of the array (3-4 weeks post-implantation), we commenced head-unrestrained datalogger recordings of extracellular activity from the 96 implanted electrodes while the animals performed the DML distractor task. A detailed description of unrestrained datalogger-based recordings is presented in Wong et al. (2023).

Neural data underwent initial processing with a common median filter to mitigate large movement-related artifacts. Subsequently, the data were further processed using a 4-pole Butterworth high-pass filter with a cutoff frequency of 500 Hz. Spike detection and sorting were then carried out offline using Plexon Offline Sorter v3. For our analysis, we included only those clearly isolated single units that exhibited average discharge rates exceeding 0.5 Hz across the entire session. We analyzed the activity of a total of 1399 units from marmoset H (13 sessions) and 1319 units (20 sessions) from marmoset D.

To ensure that animals were using mnemonic resources rather than strategic responding to perform the task, videos of all trials in all sessions were manually scored to determine whether the animal looked toward the sample during the sample-presentation period and the distractor during the distractor-presentation period. Only experimental sessions with at least 7 correct trials in each condition where the animal looked at the sample during the sample-presentation period and did not touch the screen during the delay period were included. Trials on which a distractor was presented, and the animal did not look at the screen were excluded from analysis.

### Data analysis

Analysis was performed using custom code written in Matlab (MathWorks). Statistical significance was evaluated at an alpha level of p < 0.05.

#### Task-related Activity and Spatial Tuning

For no-distractor trials, we evaluated the presence and temporal dynamics of task-related activity via two-way analyses of variance (ANOVA) conducted on mean discharge rates computed within distinct non-overlapping task epochs chosen to correspond to the various elements of the DML task. These ANOVAs included the factors location and task epoch. Location was assessed at four levels corresponding to the four spatial locations at which the visual stimuli were presented. Task epoch was assessed at six levels corresponding to the time epochs baseline, task-initiation, sample, delay, pre-response, and post-response. The baseline epoch was 3s in duration commencing 3.5 seconds prior to and ending 500ms prior to sample onset. The task-initiation epoch was 600ms in duration, beginning 500ms prior to the onset of the sample stimulus and ending 100ms after sample onset. The sample epoch was 1000ms in duration, beginning 100ms following the onset of the sample stimulus. The first 100ms following sample onset were excluded to avoid any potential task-initiation-related activity. The delay epoch was 4s in duration. The pre-response epoch was defined as the 300ms period prior to the touch response and the post-response epoch was defined as the 1000ms immediately after the touch response.

We additionally quantified the modulation of neural responses by distractor stimuli by computing a suite of contrast ratios for various epochs, which are detailed below.

#### Responses to sample and distractor stimuli

We reasoned that effective suppression of distractor interference would be reflected by an attenuation in responses to distractors relative to sample stimuli. To quantify this, we computed and compared modulation indices for sample and distractor stimuli. For sample responses, these values were calculated as a contrast ratio using mean firing rates (FR):

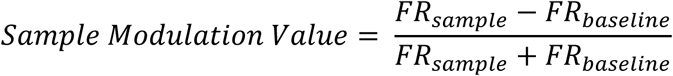

The baseline epoch was defined as the 1500 ms prior to the onset of the sample stimulus to the time of the sample stimulus presentation and the sample epoch was defined as the 100-1100 ms after the onset of the sample stimulus. Values of the index could range from –1.0 to 1.0, with positive values indicating an increase in discharge rate for the sample stimulus relative to baseline, and negative values indicating a decrease in discharge rate relative to baseline. Values of + or – 0.33 indicate a discharge rate for the sample stimulus double that of baseline.

For distractor stimuli, we computed the contrast ratio as follows:

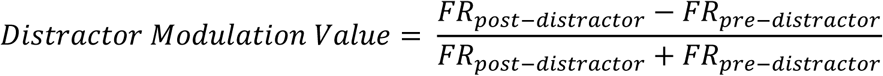

The pre-distractor epoch was defined as the 400 ms prior to the onset of the distractor to 100 ms after the distractor presentation during the delay epoch, whereas the post-distractor epoch was defined as the 100-600 ms after distractor presentation. In this case, we used as a baseline the delay period activity preceding distractor presentation, based on the logic that any visual response to the distractor would be superimposed on the activity immediately preceding it. Negative index values indicated a decrease in delay period activity following distractor onset, while positive values indicated an increase.

#### Delay-period Activity During Distractor Presentation

To investigate effects of distractor interference on spatially selective delay-period activity, we computed and compared modulation indices quantifying the dynamic range of delay period activity by comparing activity for the most versus least preferred of the four spatial locations at which samples were presented on no-distractor and distractor trials. We defined most and least preferred locations as the maximal and minimal absolute differences in discharge rates of during the delay epoch relative to baseline. Modulation indices for no-distractor trials were computed as a contrast ratio using mean firing rates (FR) during the delay epoch (1s window, 100ms after distractor presentation). The mean firing rate for the same preferred and non-preferred conditions were used to calculate the modulation indices for distractor trials.

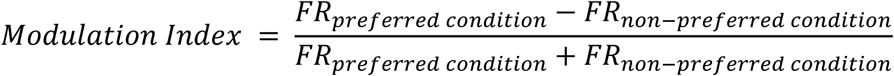

#### Relationship between neural activity and trial outcome

To investigate the relationship between distractor-induced changes in persistent activity, we computed for each spatially tuned neuron with significant delay period activity a point-biserial correlation between trial outcome and delay-period discharge rate for distractor trials. Trial outcome was coded as a binary variable (correct = 0, error = 1), such that positive coefficients indicated higher firing on error trials and negative coefficients indicated higher firing on correct trials. As with a standard Pearson correlation coefficient, the point-biserial coefficient ranges from -1 to 1. Values near 0 indicate little or no association between firing rate and trial outcome, positive values indicate greater firing on error trials, and negative values indicate greater firing on correct trials. Population-level effects were evaluated using a Wilcoxon signed-rank test against zero. Neurons were classified as showing significantly higher discharge rates on error or correct trials when their neuron-wise point-biserial coefficient was significant at p < 0.05 and positive or negative, respectively. To compare outcome-related modulation between incongruent and congruent distractor sessions, the distributions of neuron-wise point-biserial coefficients were compared using a Wilcoxon rank-sum test.

### Waveform preprocessing and cell type classification

For each single unit, the mean waveform was interpolated, using a cubic spline, from an original sampling rate of 30 kHz-1 MHz. For cell class classification, we computed two measures of the resultant waveform: trough-to-peak duration and time for repolarization. The time for repolarization was defined as the time at which the waveform amplitude decayed 25% from its peak value (Ardid et al. 2015). Previously (Wong et al. 2023) and here, we clustered cell classes into broad and narrow spiking cells using an unsupervised algorithm by Trainito et al. (2019), which correspond to putative pyramidal cells and interneurons. In this method, the expectation-maximization (EM) algorithm is used to estimate the parameters of the Gaussian mixture model (GMM), a statistical model which describes the mean and variance of underlying subpopulations. Cells with triphasic waveforms have a pronounced first positive peak followed by a larger negative trough and then a second positive peak. These cells were excluded from this analysis (n = 175), as broad and narrow spiking cells represents the most common types (Henze et al. 2000; Gold et al. 2006; Gold et al. 2009).

### Array localization

At the end of data acquisition, Marmosets were humanely euthanized and brains perfused in order to allow reconstruction of recording sites (Wong et al. 2023). The animals were deeply anesthetized with 20 mg/kg of ketamine plus 0.025 mg/kg medetomidine and 5% isoflurane in 1.4–2% oxygen to reach beyond the surgical plane (i.e., no response to toe pinching or cornea touching). They were then transcardially perfused with 0.9% sodium chloride irrigation solution, followed by 10% buffered formalin. The brain was then extracted and stored in 10% buffered formalin for more than a week before ex-vivo magnetic resonance imaging (MRI). On the day of the MRI scan, the brain was transferred to a 50 mL conical centrifuge tube and immersed in a fluorine-based lubricant (Christo-lube; Lubrication Technology) to improve homogeneity and avoid susceptibility artifacts at the boundaries. Ex-vivo MRI was performed on a 15.2T 11-cm horizontal bore magnet (Bruker BioSpin Corp, Billerica, MA) and Bruker BioSpec Avance Neo console with the software package Paravision 360.3.5 (Bruker BioSpin Corp, Billerica, MA). The Bruker gradient/shim coil (B-GA6S-100) had a 6-cm inner diameter, a maximum gradient strength and slew rate of 1000 mT/m and 9000 T/m/s, respectively, and a full set of 3^rd^-order shims. The radiofrequency coil was a 35-mm inner diameter, transmit/receive volume coil with quadrature detection (Bruker BioSpin Corp, Billerica, MA). High resolution (70 µm isometric) T1-flash images were acquired for each animal with the following parameters: resolution time (TR) = 0.026s, echo time (TE) = 0.008s, slice thickness = 0.07mm, flip angle = 20. The raw MRI images were converted to NIfTI format using dcm2niix (Li et al. 2016), and then manually registered to the high-resolution ex vivo NIH template brain (Liu et al. 2018). The boundaries of the Paxinos atlas (Paxinos et al. 2012), included in the NIH template, were overlaid on the registered ex vivo anatomical T2 images, were used to reconstruct the location of the implanted array in each marmoset. These cytoarchitectonic boundaries overlayed on the registered ex-vivo anatomical T2 images were used to reconstruct the location of the implanted array in each marmoset. Images were rendered in 3-D using the program *MRIcroGL* and overlayed the Paxinos atlas cytoarchitectural boundaries for estimating where the arrays were implanted in each animal (Figure 2).

**Figure 2.**
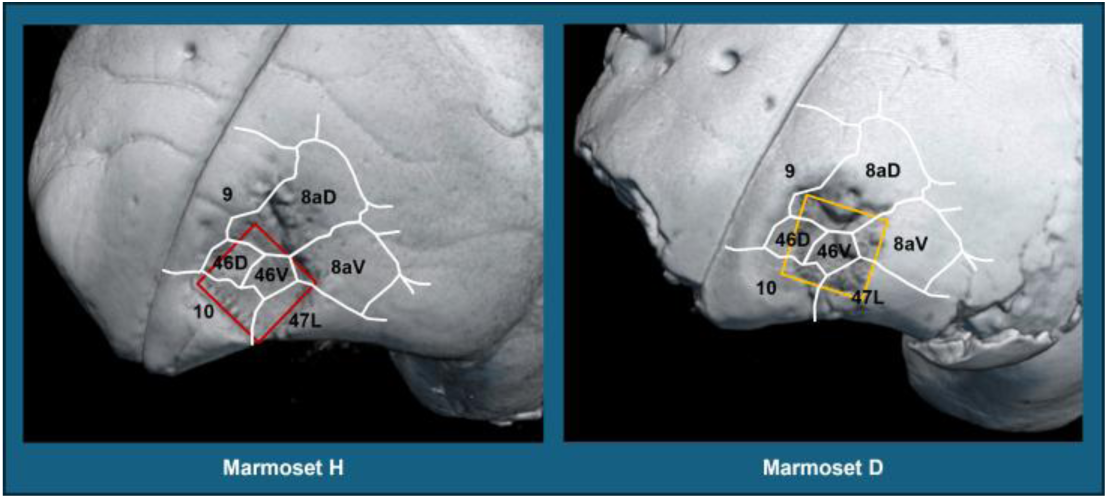
Reconstruction of recording sites. T1-flash MRIs were acquired and manually registered to the NIH template brain that contains the location of cytoarchitectonic boundaries of the Paxinos atlas (see methods). Images were rendered in 3-D using *MRIcroGL* and overlayed with the Paxinos atlas cytoarchitectural boundaries on: Marmoset Brain Template (Left panel); Marmoset H (Middle panel); Marmoset D (Right panel). Arrays in both animals largely covered areas 46D and 46V, with reduced coverage of areas 8aD, 8aV, 10, and 47.

## Results

### Task performance

Marmosets participated in a total of 33 (17 incongruent and 18 congruent) sessions in which single-unit activity in PFC was recorded during performance of the DML task. Overall, marmosets were able to perform the DML task with a 4s delay period; accuracy level were above chance (25%) for both distractor and no-distractor conditions (Figure 1B). We additionally observed an interaction between distractor condition and distractor type on task performance. On incongruent distractor sessions, the task accuracies between no distractor (x̅ = 74.3%) and distractor trials (x̅ = 75.1%) were not significantly different (*P* = 0.70). In contrast, on congruent sessions, task accuracy was reduced by distractor presentation (no-distractor trials, x̅ = 70.6%, distractor trials, x̅ = 66.0%, P <0.05). Contrasting session types, more error responses were directed towards the distractor location in congruent (54.2%) than incongruent (37.7%) sessions (Figure 1C).

To further investigate task performance, we compared marmosets’ reaction times (RTs) on correct and error trials at the 4s delay duration for both session types. If animals were relying on mnemonic processes to guide response selection, we reasoned that RTs on error trials would be similar to or longer than those on correct trials, reflecting “diligent guesses” regarding the remembered sample location (Link 1982). We found that RTs were significantly greater for error than correct trials in both distractor conditions and session types (*t*-test, *P* < 0.05) (Incongruent: no distractor x̅ correct RT = 990ms, no distractor x̅ error RT = 1220ms, distractor x̅ correct RT = 940ms, distractor x̅ error RT = 1170ms; Congruent: no distractor x̅ correct RT = 980ms, no distractor x̅ error RT = 1250ms, distractor x̅ correct RT = 990ms, distractor x̅ error RT = 1310ms). These results are consistent with a reliance of marmosets on mnemonic processes during DML task performance and indicate that performance on the DML task was an accurate reflection of marmosets’ spatial WM abilities.

### Marmoset PFC neurons exhibit task-related activity

We recorded the activity of a total of 2718 PFC neurons (1399 in Marmoset H, 1319 in Marmoset D) in two monkeys over the 33 experimental sessions in which they performed the DML task. We defined task-related modulation as a significant difference in mean discharge rate between any task epoch and the baseline period, as evaluated using ANOVA. Many neurons exhibited task-related activity modulations, and these modulations were observed in all epochs of the DML task. Representative single neurons exhibiting activity modulations during the task-initiation, sample, delay, and response epochs are shown in Figure 3. A majority of neurons showed significant modulations in discharge rates for both distractor types (incongruent: 1111/1212 (91.7%), congruent: 1367/1506 (90.7%)). Of the 1111 neurons modulated during incongruent distractor sessions, 275 (22.7% of overall neurons) displayed task-initiation-related activity, 665 (54.9%) displayed sample-related activity, 736 (60.7%) displayed delay-period activity, 547 (45.1%) displayed pre-response-related activity, and 685 (56.5%) displayed post-response-related activity. Of the 1367 neurons modulated during congruent colour distractor sessions, 298 (19.8% of overall neurons) displayed task-initiation-related activity, 749 (49.7%) displayed sample-related activity, 972 (64.5%) displayed delay-period activity, 536 (35.6%) displayed pre-response-related activity, and 764 (50.7%) displayed post-response related activity. Overall, we found that, across both animals, well-isolated single units were recorded across a broad set of prefrontal subregions including areas 46D, 46V, 8aD, 8aV, 47 and 10 (see Figure 4). We observed task-related activity in all epochs and in all subregions.

**Figure 3.**
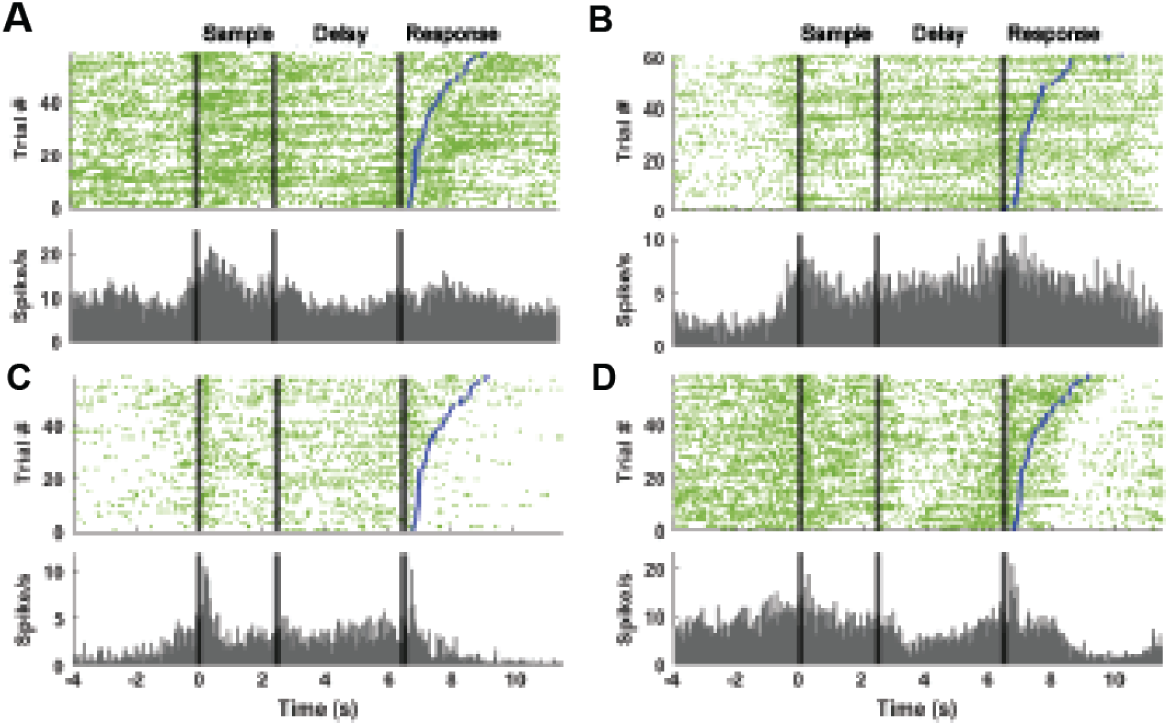
Example PFC neurons exhibiting sample- and delay-period activity during the DML task. Relative to baseline, neurons in **(A-C)** showed increased firing rate during both the sample-and delay periods, whereas the neuron in (**D**) showed increased firing rate during the sample period, but decreased firing rate during the delay-period. Rasters are aligned to sample-onset. Blue lines indicate reaction time.

**Figure 4.**
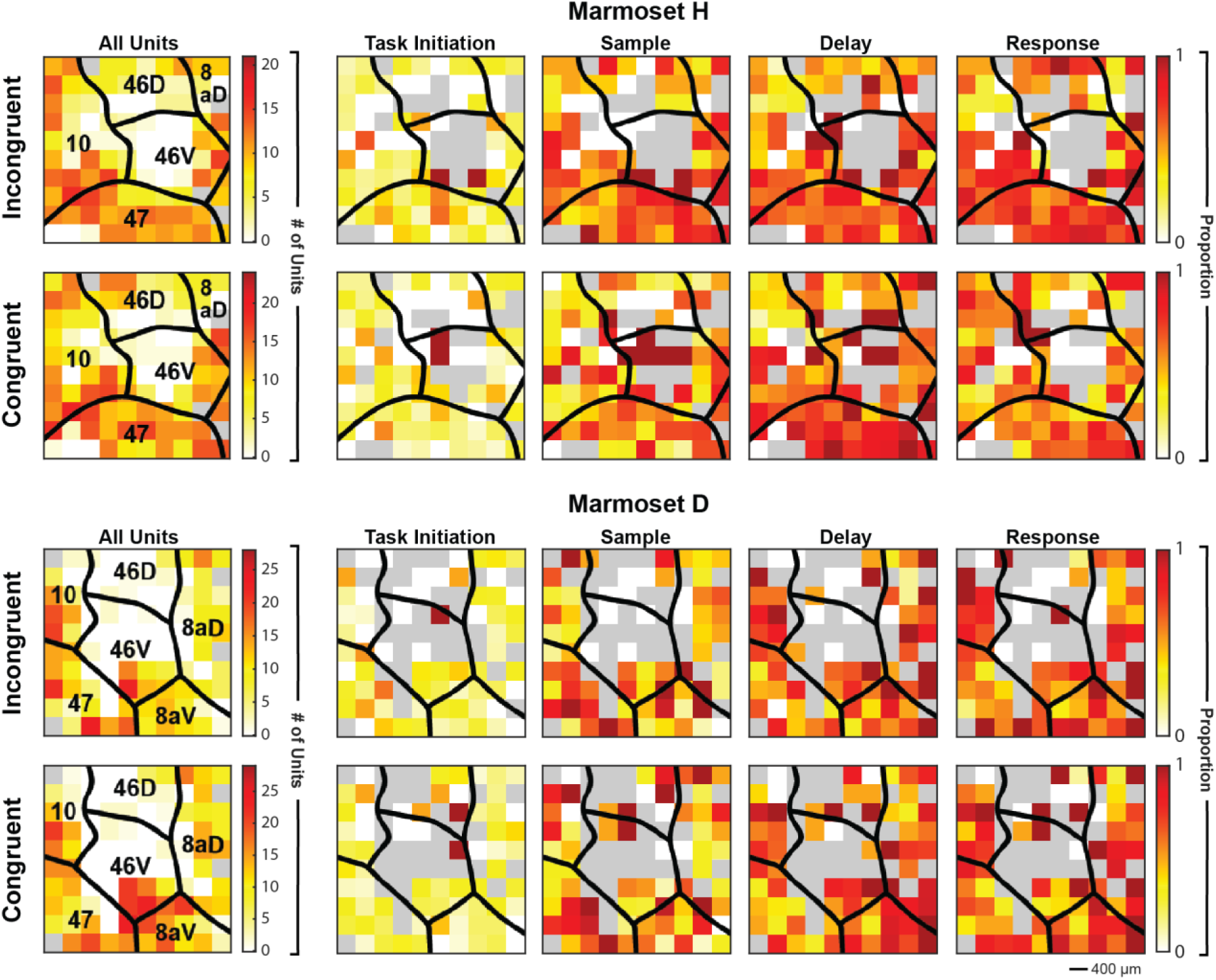
Distribution of task-modulated units. Array locations were reconstructed using high-resolution MRIs and superimposed on a standardized marmoset brain, area boundaries from Paxinos et al. (2012). The first column (left column) represents the total number of units found across sessions and its distribution on the array. The second to fifth column represents the proportion of units compared to the first column. For each marmoset and in both distractor colour session types, task-modulated units were distributed across the array in locations where well-isolated single units were observed. Grey colour depicts array locations at which well-isolated single units were not observed.

Previous work in macaques performing delayed-response tasks has revealed that modulations of sample-, delay- and response-related activity may take the form of either increases or decreases from the baseline discharge rate (Funahashi et al. 1989; Funahashi et al. 1991). To investigate such modulations in marmoset PFC, we classified each significantly task-modulated neuron as excited or suppressed relative to baseline within each task epoch (Table 1) and then tested whether the distribution of modulation sign differed across epochs using chi-squared tests on epoch × modulation-sign contingency tables, separately for congruent and incongruent conditions. The distribution of excitation and suppression varied significantly across epochs in both incongruent (X^2^ (4, *N* = 2908) = 149.98, *P* < 0.05) and congruent sessions (X^2^ (4, *N* = 3319) = 176.71, *P* < 0.05). In both session types, excitation was more prevalent overall; however, the proportion of suppressed neurons increased during the delay epoch relative to several other epochs. Specifically, suppression reached 32.6% in incongruent sessions and 48.4% in congruent sessions during the delay, representing the highest proportion observed across epochs in the congruent condition. These results show that the pattern of task-related modulation changed significantly across epochs, with the delay period associated with the greatest relative increase in suppression.

**Table 1.**
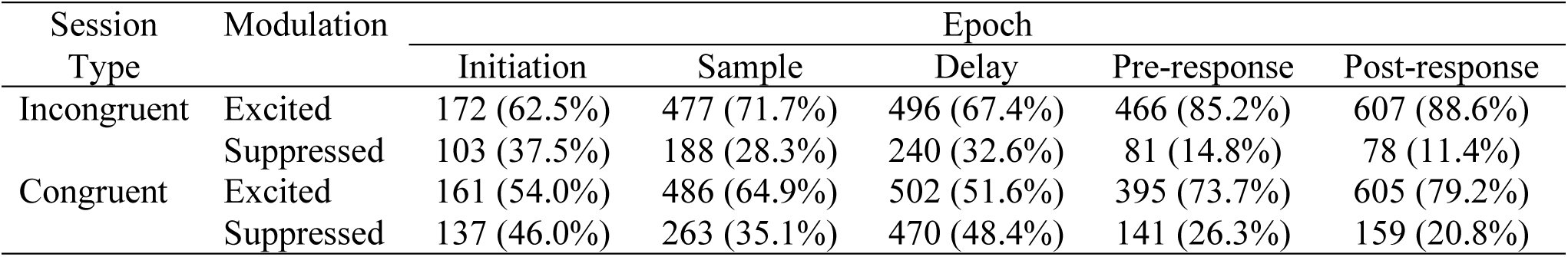
Number of neurons excited or suppressed in each task epoch for incongruent and congruent sessions.

| Session Type | Modulation | Epoch |  |  |  |  |
| --- | --- | --- | --- | --- | --- | --- |
|  |  | Initiation | Sample | Delay | Pre-response | Post-response |
| Incongruent | Excited | 172 (62.5%) | 477 (71.7%) | 496 (67.4%) | 466 (85.2%) | 607 (88.6%) |
|  | Suppressed | 103 (37.5%) | 188 (28.3%) | 240 (32.6%) | 81 (14.8%) | 78 (11.4%) |
| Congruent | Excited | 161 (54.0%) | 486 (64.9%) | 502 (51.6%) | 395 (73.7%) | 605 (79.2%) |
|  | Suppressed | 137 (46.0%) | 263 (35.1%) | 470 (48.4%) | 141 (26.3%) | 159 (20.8%) |

### Marmoset PFC neurons exhibit spatial tuning during DML task performance

In marmoset PFC, we observed single units that exhibited spatial tuning of discharge rates, relative to baseline activity, during the sample, delay and response epochs of the DML task. Spatial tuning was assessed using one-way ANOVAs to compare the mean discharge rates of each neuron across conditions (four spatial locations) and each task epochs relative to baseline. Neurons showing a significant interaction and significant post-hoc comparisons were classified as spatially tuned. In incongruent sessions, we found 354 neurons (53.2% of modulated neurons) that exhibited tuning during the sample period, 295 neurons (40.1%) displayed tuning during the delay-period, 155 neurons (28.3%) displayed tuning during the pre-response period, and 174 neurons (25.4%) displayed tuning during the post-response period. In congruent sessions, 372 (49.7%), 382 (39.3%), 169 (31.5%) and 224 (29.3%) neurons displayed tuning activity during the sample period, delay-period, pre-response period and post-response period, respectively.

### Distractor response is reduced relative to sample

To investigate potential distractor filtering in PFC persistent activity, we calculated separate modulation indices based on discharge rates relative to baseline activity during sample and distractor presentations (see methods). Average sample and distractor modulation values were computed separately for neurons that were classified as sample-related, delay-related or both. In incongruent colour sessions, the mean sample modulation value was 0.1119 and the mean distractor modulation value was 0.1601 (n = 976). In congruent sessions, the mean sample and distractor modulation values were 0.1200 and 0.1509, respectively (n = 1202). In both session types, the mean sample modulation value was greater than the distractor modulation value (*P* < 0.001), indicating that the response to the distractor during the delay period was attenuated relative to the response to the sample (Figure 5).

**Figure 5.**
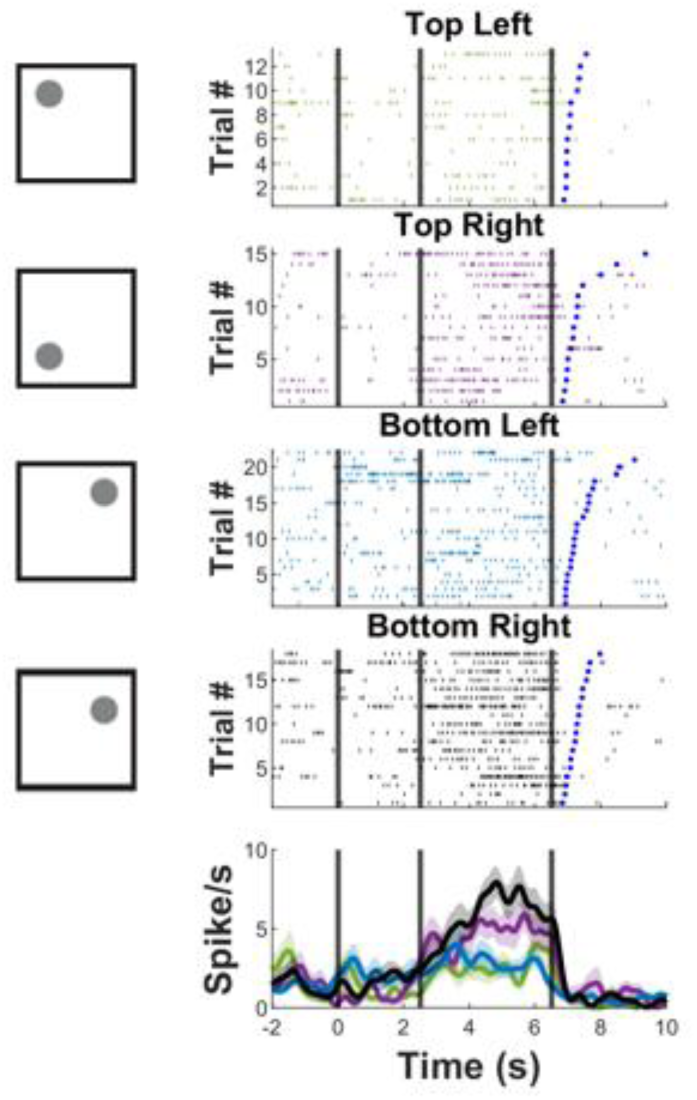
Representative PFC neuron exhibiting spatial tuning during the DML task, with selective responses to the bottom-right location. Bottom panel displays the spike density function (mean ± SEM). Rasters are aligned to sample-onset, and the blue lines indicate reaction time.

### Delay-period activity is modulated by distractor presentation

To investigate whether distractor presentation disrupted spatial tuning, we computed a modulation index for spatially tuned delay neurons comparing preferred and non-preferred conditions separately for distractor and no-distractor trials (Figure 6). On incongruent colour sessions, the mean modulation index for no distractor trials was 0.2749 and -0.0031 for distractor trials (n = 295). In congruent sessions, the modulation indices for no distractor and distractor trials are 0.2890 and -0.0055 (n = 382), respectively. In both session types, the modulation index for distractor trials was significantly lower than that on no distractor trials (*P* < 0.001), indicating that the difference in activity between preferred and non-preferred locations was reduced on distractor trials. Overall, this pattern of results indicates that spatial tuning, was abolished or weakened in response to distractor presentation.

**Figure 6.**
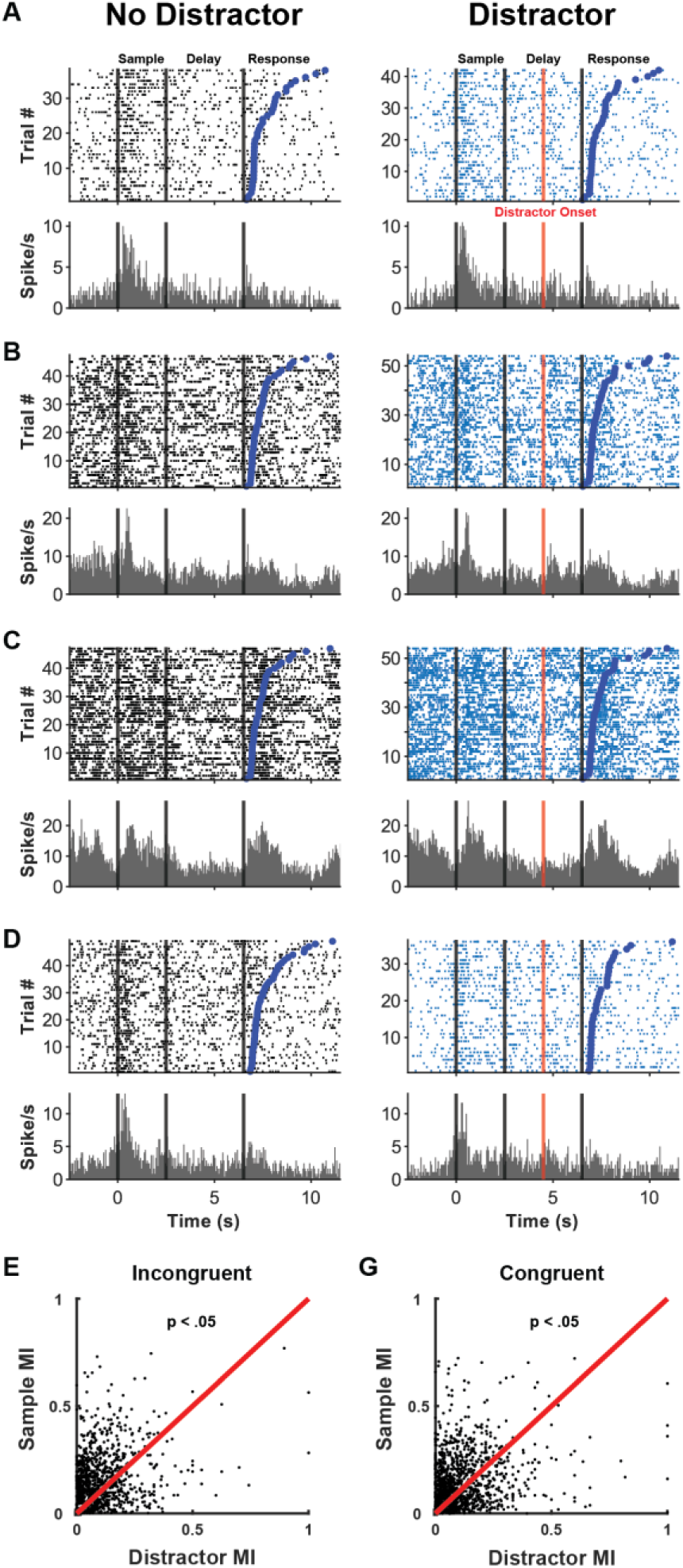
PFC neurons exhibit suppressed responses to distractors relative to sample stimuli. Rasters and peristimulus time histograms for example neurons comparing no-distractor (black, left panels) and distractor trials (blue, right panels) (**A-E**). Lower panels, scatterplots comparing modulation indices for sample and distractor stimuli. Modulation indices were calculated as a contrast ratio using mean firing rates (see Methods) and plotted against each other for incongruent colour distractor sessions (**F**) and congruent colour distractor sessions (**G**).

### Persistent delay-period activity in marmoset PFC reflects task performance

Previous work has shown that persistent delay-period activity is attenuated on error trials relative to correct trials (Funahashi et al. 1989). To investigate this in marmoset PFC, we compared the discharge rates between correct and error trials (no distractor trials) for neurons which were responsive during the delay period (e.g. exhibited discharge rates significantly different during the delay epoch relative to baseline). Overall, 366/677 (54.1%; independent samples *t*-test, *P* < 0.05) neurons had a significant difference in discharge rate between correct and error trials during the delay period (incongruent: 159/295 (53.9%); congruent: 207/382 (54.2%). Figure 7A depicts an example neuron for which the discharge rate during the delay period was significantly different between correct and error trials.

**Figure 7.**
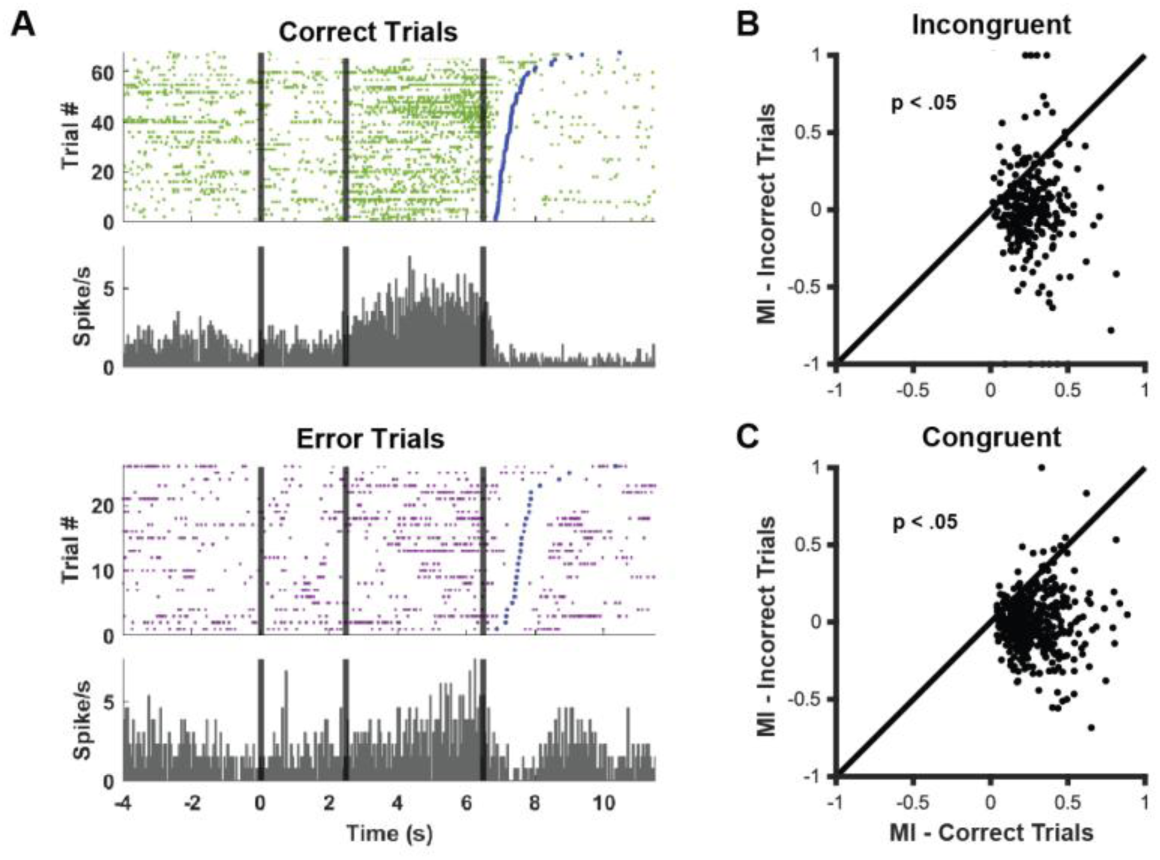
Delay-period activity reflects task performance. Example neuron for which the discharge rate during the delay-period differed significantly between correct and error trials **(A).** Delay-period modulation index computed from the preferred and non-preferred conditions separately for correct and error trials **(B-C).** Overall, delay-period activity is reduced on error trials.

To further investigate the magnitude of differences in neural activity between correct and error trials during the delay period, we computed a modulation index from the preferred and non-preferred conditions separately for correct and error trials (Figure 7B-C). For incongruent sessions, the modulation index was higher for correct trials than error trials (0.3303 vs. 0.0730, n = 295, *P* < 0.001). This trend was similar for congruent sessions (0.3567 vs. 0.1269, n= 382, *P* < 0.001). Overall, these data are consistent with previous reports indicating that delay-period activity was reduced on error trials.

### Localization of Spatially Selective Delay Activity

To determine the PFC locations at which we observed the strongest, spatially selective delay activity, we averaged the modulation indices of neurons recorded at each electrode contact of the Utah array across sessions. We did not observe any spatial organization of modulation index across these arrays (Figure 8).

**Figure 8.**
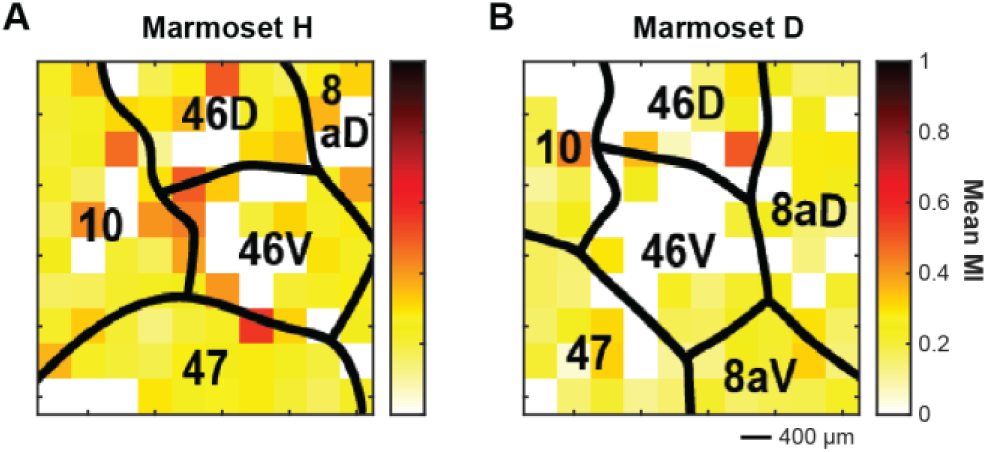
Location of spatially selective delay activity. Average modulation indices of neurons recorded at each electrode contact of the Utah array across sessions for Marmoset H **(A)** and Marmoset D **(B)**. No particular region of the array exhibited a consistently higher average modulation index than the other regions.

### Task performance is correlated with distractor interference

To determine whether task performance was correlated with distractor interference, we examined delay-period activity in spatially tuned delay neurons (n = 677) as a function of distractor condition and trial outcome. We first compared firing rates on distractor trials between correct and error outcomes (ANOVA). Of the 295 spatially tuned delay neurons recorded during incongruent colour distractor sessions, 37 neurons (12.5%) showed a significant difference between correct and error distractor trials (p < 0.05). Of these, 33 neurons (89.2%) had higher discharge rates on error trials, whereas 4 neurons (10.8%) had higher discharge rates on correct trials. In congruent colour distractor sessions, 96 of 382 spatially tuned delay neurons (25.1%) showed a significant difference between correct and error distractor trials. Of these, 88 neurons (91.7%) had higher discharge rates on error trials, whereas 8 neurons (8.3%) had higher discharge rates on correct trials. Representative neurons are shown in Figure 9A; Figure 9B summarizes firing rates for individual neurons, and Figure 9C shows the population mean firing rate.

**Figure 9.**
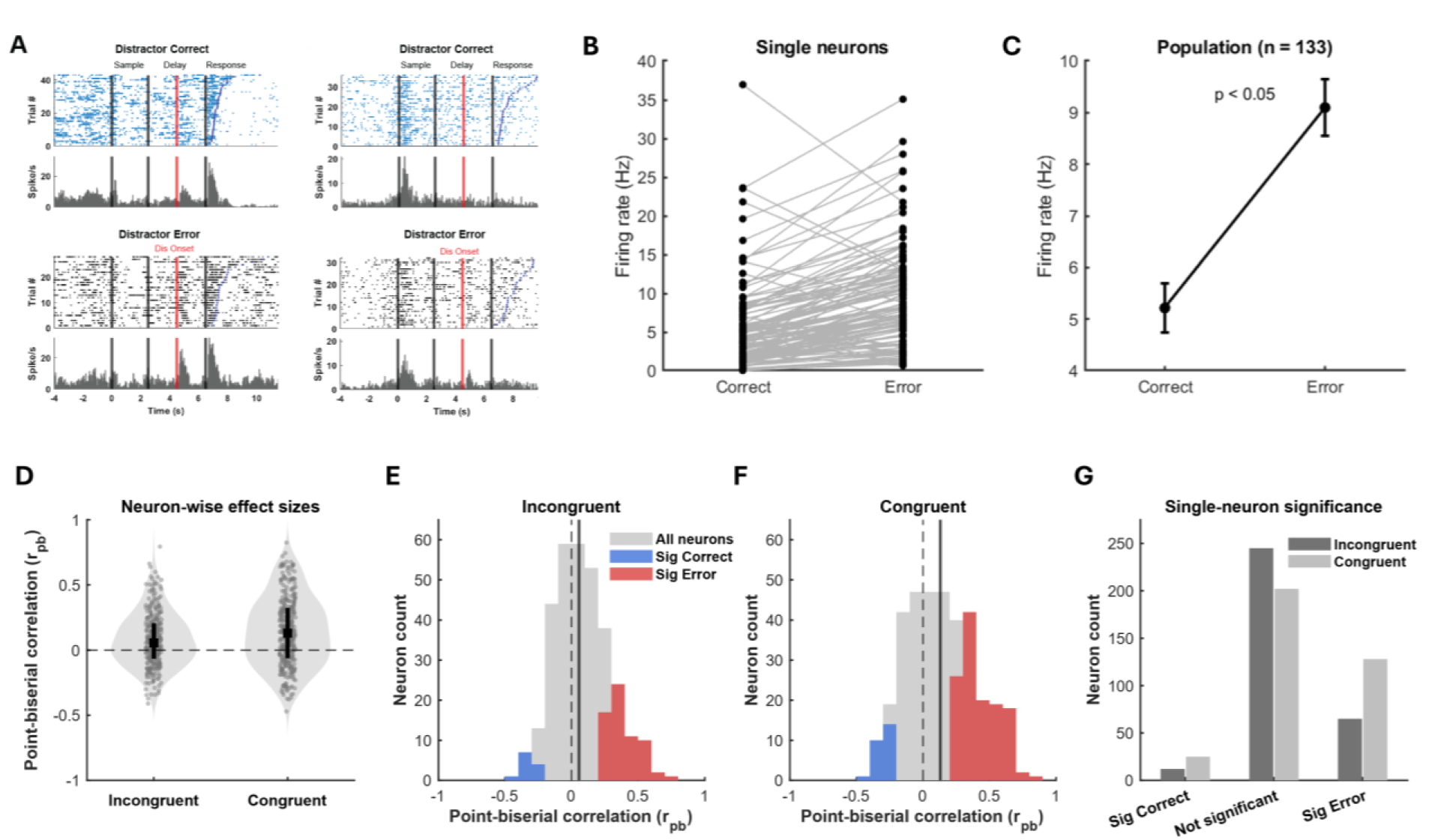
Task performance is associated with distractor interference. **(A)** Representative delay-period spatially tuned neurons showing higher discharge rates on distractor error trials than on distractor correct trials. Raster plots and PSTH are aligned to sample onset; distractor onset is indicated by the red line. **(B)** Summary of delay-period firing rates for individual neurons on distractor correct and distractor error trials. Each line connects the responses of a single neuron across the two trial outcomes. **(C)** Population mean firing rate across neurons shown in panel B. Error bars indicate SEM. **(D)** Neuron-wise point-biserial correlation coefficients (r_pb) relating delay-period firing rate to trial outcome on distractor trials across all spatially tuned delay neurons. Positive coefficients indicate stronger delay activity on error trials, whereas negative coefficients indicate stronger delay activity on correct trials. Violin plots show the distribution across neurons, with black symbols indicating the median and interquartile range. **(E)** Distribution of neuron-wise point-biserial coefficients in incongruent distractor sessions. Gray indicates all neurons, blue indicates neurons with significantly higher discharge rates on correct trials, and red indicates neurons with significantly higher discharge rates on error trials. **(F)** Distribution of neuron-wise point-biserial coefficients in congruent distractor sessions, plotted as in panel E. **(G)** Number of neurons showing significantly higher discharge rates on correct trials, no significant outcome-related difference, or significantly higher discharge rates on error trials in incongruent and congruent distractor sessions.

We next quantified this relationship at the population level using a point-biserial correlation analysis restricted to distractor trials, where positive coefficients reflect stronger delay activity on error trials and negative coefficients reflect stronger delay activity on correct trials. Across all 677 spatially tuned delay neurons, point-biserial coefficients were significantly shifted above zero (r̄_pb_ = 0.112, r̄_pb_ = 0.090, signed-rank test, p < 0.05), indicating that delay-period firing was generally higher on distractor error than correct trials (Figure 9D). When analysed separately by session type, this positive shift was present in both incongruent and congruent sessions but was significantly larger in congruent sessions. In incongruent sessions (n = 322), coefficients were significantly greater than zero (r̄_pb_ = 0.080, r̄_pb_ = 0.056, signed-rank test, p < 0.05; Figure 9E), with 65 neurons (20.2%) exhibiting significantly higher activity on error trials and far fewer (12 neurons - 3.7%) exhibiting significantly higher activity for correct trials (Figure 9G). In congruent sessions (n = 355), this effect was larger (r̄_pb_ = 0.141, r̄_pb_ = 0.130, p < 0.05; Figure 9F), with 128 neurons (36.1%) exhibiting significantly greater error trial discharge rates, and 25 neurons (7.0%) with greater activity on correct trials. (Figure 9G). The two distributions differed significantly from one another (rank-sum test, p = 0.0036).

Together, these results indicate that distractor-related errors were associated with elevated delay-period activity in spatially tuned neurons, and that this relationship was stronger during congruent than incongruent distractor sessions. These findings are consistent with the behavioural reduction in task performance observed on congruent, but not incongruent, distractor trials and support an association between distractor filtering at the single neuron level and task performance.

### Broad and narrow spiking neurons are modulated in all task epochs

Broad spiking (BS) and narrow spiking (NS) cells have previously been demonstrated to contribute differentially to spatial tuning in the oculomotor delayed-response task (Rao et al. 1999). To investigate the contribution of putative pyramidal cells and interneurons to mnemonic processes in marmoset PFC, we used the EM algorithm for GMM clustering method on the trough-to-peak duration and time for repolarization of single unit waveforms to identify BS and NS cell clusters in an unsupervised manner. This resulted in 2045/2543 cells (80.4%) being classified as broad spiking and 498/2543 cells (19.6%) being classified as narrow spiking cells. For neurons modulated in each epoch of the DML task, we identified whether the neuron was classified as BS or NS. 434 BS (75.7%) and 102 NS (17.8%) cells displayed task-initiation-related activity; 1033 BS (73.1%) and 289 NS (20.4%) cells displayed sample-related activity; 1285 BS (75.2%) and 322 NS (18.9%) cells displayed delay-period activity; 813 BS (75.1%) and 216 NS (19.9%) cells displayed pre-response-related activity; and 1070 BS (73.8%) and 307 NS (21.2%) cells displayed post-response-related activity. For the task-modulated cells, a chi-square goodness of fit test was conducted to determine if the proportion of each cell type varied across the task epochs. No significant differences were observed (X^2^ (4, *N* = 5871) = 4.117, *P* > 0.05). We further identified whether BS and NS cells were excited or suppressed in each task epoch compared to baseline (Table 2). Overall, consistent with previous reports in marmoset (Wong et al. 2023) and macaque PFC (Wilson et al. 1994; Rao et al. 1999; Constantinidis and Goldman-Rakic 2002), we observed that both BS and NS neurons were modulated in all task epochs and that this modulation could take the form of excitation or suppression.

**Table 2.** Number of neurons excited or suppressed in each task epoch separated by cell type.

| Modulation | Cell Type | Epoch |  |  |  |  |
| --- | --- | --- | --- | --- | --- | --- |
|  |  | Initiation | Sample | Delay | Pre-response | Post-response |
| Excited | BS | 243 | 702 | 760 | 639 | 895 |
|  | NS | 67 | 210 | 187 | 186 | 269 |
| Suppressed | BS | 191 | 331 | 525 | 174 | 175 |
|  | NS | 35 | 79 | 135 | 30 | 38 |

## Discussion

Here, we used a touchscreen-based DML task to investigate whether marmoset WM performance was robust to distractor interference, the effects of salient visual distractors on PFC persistent activity, and the link between distractor-mediated modulations in persistent activity and behavioural performance. We found that marmosets’ overall performance on this task was robust to distractor interference. On congruent distractor trials, in which colour of the distractor matched that of the sample stimulus, marmosets exhibited only a modest reduction in task performance, while incongruent colour distractors had no effect on performance. These behavioural effects were mirrored by the effects of distractors on PFC persistent activity in two respects. First, while distractor presentation did modulate persistent activity, we found that neural responses to distractors were significantly attenuated relative to responses evoked by sample stimuli, suggesting an active process of distractor suppression in support of WM performance. Second, we found that distractor-related activity was greater on error than correct trials, and this was evident within a much larger proportion of PFC neurons for congruent than incongruent distractors. This suggests a failure of active suppression on error trials which is more prevalent for congruent distractors. These findings establish marmosets as a viable model for investigating the behavioural and neural basis of distractor filtering during WM and provide evidence for conserved mechanisms of interference control in primate PFC.

Our findings are consistent with prior work in rhesus macaques demonstrating a critical role for dlPFC in distractor suppression (Miller et al. 1996; Everling et al. 2006). Similar to the macaque dlPFC (Suzuki and Gottlieb 2013), marmoset PFC neurons responded weakly to salient distractors presented during the delay period, with responses attenuated relative to the sample stimulus. Critically, we replicated the key finding that distractor responses were reduced on correct trials relative to error trials, indicating a link between effective distractor suppression and successful WM performance. This parallel extends to the behavioral level: reversible inactivation of macaque dlPFC produces substantial impairments in distractor suppression (Suzuki and Gottlieb 2013), and our data show that marmosets with intact PFC can successfully filter distractors, though performance is degraded when distractor salience increases. The correspondence between marmoset and macaque PFC in both neural response patterns and their relationship to behavior supports the hypothesis that distractor filtering mechanisms are conserved across primate species and validates the marmoset as a model for investigating these processes.

In line with our previous findings (Wong et al. 2023; Wong et al. 2024), we found that marmoset PFC neurons exhibited sample-, delay-, and response-related activity during the DML task with response profiles similar to those observed in the earliest reports of persistent activity in macaque PFC (Fuster and Alexander 1971; Kubota and Niki 1971). As noted there, a distinguishing feature of delay cells was spatial tuning for the cued location (Funahashi et al. 1989; Funahashi et al. 1990; Funahashi 2006). We additionally observed that delay-period activity was significantly stronger on correct trials than on error trials during no-distractor conditions, consistent with the classic finding that persistent activity in macaque dlPFC is diminished or truncated on error trials (Funahashi et al. 1989). In contrast, during distractor sessions, delay-period activity of spatially tuned neurons was generally elevated on error relative to correct trials, as shown by both the single-neuron ANOVA results and the positive shift in point-biserial coefficients across the population. This relationship was present in both incongruent and congruent distractor sessions, but was significantly stronger in congruent sessions, paralleling the greater behavioural impairment observed under that condition. This interpretation is broadly consistent with evidence that macaque dlPFC supports resistance to distraction during memory-guided behaviour, with distractor-related activity being linked to subsequent errors when interference is not successfully suppressed (Suzuki and Gottlieb 2013).

Despite these similarities, we observed that persistent delay-period activity in marmosets appeared weaker and less spatially selective than typically observed in macaques performing oculomotor delayed response (ODR) tasks. This is most likely attributable to the nature of the DML task we employed as a measure of WM here. We elected to use this paradigm in freely moving animals on the basis of the fact that, consistent with observations of high visual distractibility (Magrou et al. 2026), marmosets perform relatively poorly on the classic ODR task under head restraint, with accuracy well below that of rhesus macaques even at short delays of 100-400ms (Amly et al. 2025). This design difference results in at least two fundamental changes in task demands which may have influenced the persistent activity we observed. First, it shifts the potential reference frame in which marmosets were working to allocentric rather than retinocentric as in the ODR task. Human fMRI studies have shown that egocentric representations often elicit greater activation than allocentric representations during working memory tasks (Moraresku and Vlcek 2020), which is consistent with the reduction in the magnitude of persistent activity we observed. Second, performance of the DML task did not require the same degree of fine spatial precision of either mnemonic representations nor motor responses as most variants of the ODR task (Funahashi et al. 1989). This is broadly consistent with the reduction in spatial tuning we observed here. It is worth noting that the combination of behavioural control and precision combined with neural recordings in studies using the ODR task has been the bedrock of much of our understanding of the mechanisms of spatial tuning in WM (Wang 2021). We suggest that the differences in persistent activity we observed here differ from those foundational observations in degree rather than kind and while they must be acknowledged, they do not discount the potential value of the marmoset model for investigations of the neural basis of WM. Indeed, combining insights from head restrained and freely moving experimental paradigms in different species has the potential to substantially broaden the scope of our understanding of WM mechanisms not only across species but also to more ecologically diverse settings. An emerging body of work in the marmoset model has aimed to combine behavioural analysis of complex naturalistic behavior with neurophysiological recordings (Miller et al. 2022; Ngo et al. 2022; Mitchell et al. 2024; Li et al. 2026). In this sense, the complementary value of the marmoset model may be to bridge the gap between the substantial mechanistic insights made possible by head restrained recordings in controlled laboratory paradigms, and the operation of those mechanisms in fluid and dynamic environments.

Our findings align with computational models proposing that dlPFC circuits instantiating WM rely on attractor networks which generate persistent activity via local recurrent excitation balanced by global inhibition (Wang et al. 2004; Wang 2009; Wang 2021). Such models make specific predictions about distractor interference. First, they predict a similarity effect, whereby distractors that are more similar to the sample stimulus generate stronger interference by activating overlapping neural populations (Furman and Wang 2008). Our behavioral data were consistent with this prediction: we found that congruent color distractors significantly impaired performance, whereas incongruent distractors did not. Second, attractor models predict that distractor suppression is critical for protecting the integrity of memory traces, with bottom-up stimulus input requiring active suppression in order to maintain stable WM representations (Compte et al. 2000; Brunel and Wang 2001). Here we found that distractor responses were attenuated relative to the sample, and that a failure to suppress distractor responses—as evidenced by stronger distractor responses on error trials—resulted in impaired performance. These results provide additional empirical support for attractor-based models of working memory and suggest that the computational principles governing distractor filtering are implemented in marmoset PFC circuits.

The mechanisms by which dlPFC reduces distractors remain an area of active investigation. Work in macaques suggests that dlPFC implements anticipatory suppression locally, reducing visual responses to irrelevant distractors without requiring inhibition of remote structures (Ruff and Driver 2006; Suzuki and Gottlieb 2013). However, anatomical evidence indicates that frontal projections to visual areas such as V4 and lateral intraparietal cortex (LIP) mainly terminate on excitatory neurons (Anderson et al. 2011), suggesting that dlPFC likely influences distractor processing through feedback enhancement of target representations rather than direct inhibition of distractor representations (Moore and Armstrong 2003). In marmosets, area 46 projects to multiple visual areas with a concentration in the dorsomedial subdivision of the dorsal stream (Burman et al. 2006), providing anatomical pathways through which marmoset PFC could modulate visual processing. Comparisons between dlPFC and LIP in macaques have shown that while both areas contribute to visual selection, LIP reflects covert shifts of attention between target and distractor locations, whereas dlPFC has a determining influence on how stimuli guide action (Bisley and Goldberg 2003; Bisley and Goldberg 2006; Suzuki and Gottlieb 2013). Future work examining the coordination between marmoset PFC and posterior visual areas during distractor filtering will be important for understanding the circuit-level mechanisms of interference control.

In both animals, we observed robust activation of area 10 (frontopolar cortex) during the sample, delay, and response epochs. This is notable because single-neuron recordings in macaque area 10 have generally not revealed persistent delay-period activity (Tsujimoto et al. 2010; Tsujimoto et al. 2012; Boschin and Buckley 2015), despite neuroimaging evidence in humans showing frontopolar cortex activation during working memory and episodic memory tasks (S. J. Gilbert et al. 2006; Sam J. Gilbert et al. 2006; Okuda et al. 2007; Burgess et al. 2011; Volle et al. 2011). In macaques, lesions of area 10 do not impair working memory per se, but rather enhance behavioral adaptation following conflict and reduce susceptibility to distractors regardless of salience (Mansouri et al. 2015; Mansouri et al. 2017). Area 10 has been hypothesized to play a unique role in redistributing executive control resources among potential goals in complex, changing situations (Boorman et al. 2009; Boschin and Buckley 2015; Mansouri et al. 2015). Combined with the relative cytoarchitectonic uniformity of marmoset area 10, anatomical tracing studies support broad homology between marmoset and macaque area 10, with major frontal, temporal, and subcortical connections that are largely similar across species (Burman et al. 2006; Roberts et al. 2007; Burman and Rosa 2009). Nevertheless, the marmoset frontal pole may exhibit somewhat broader or distinct connectional features, including possible medial–lateral subdivisions characterized by different balances of input from different sources (Burman, Reser, Yu, et al. 2011; Burman, Reser, Richardson, et al. 2011). We observed sample-, delay-, and response-related activity in marmoset area 10 during the DML task (Wong et al. 2024). However, additional animals and direct comparisons across task types will be needed to clarify its precise role and to determine whether marmoset area 10 is functionally distinct from macaque area 10. The accessibility of marmoset area 10, which lies on the cortical surface, makes this species particularly well suited for future investigations of frontopolar function.

The lissencephalic architecture of marmoset PFC, which renders the entire prefrontal cortex accessible to laminar recordings, provides a unique opportunity to investigate the microcircuit mechanisms underlying working memory and distractor filtering. Current computational models make layer- and cell-type-specific predictions about how persistent activity is generated and maintained, and how distractor interference propagates through cortical circuits (Wang et al. 2004; Wang 2021; Joyce et al. 2025). For example, dopaminergic projections from the ventral tegmental area innervate primate PFC with layer-specific density, preferentially targeting superficial layers where dopamine modulates persistent activity and distractor resistance (Williams and Goldman-Rakic 1995; Goldman-Rakic 1996; Vijayraghavan et al. 2007). Recent comparative work has revealed that marmoset dlPFC parvalbumin neurons express higher levels of dopamine D1 receptors than macaques (Joyce et al. 2025), which may influence the dynamics of distractor filtering in layer-specific microcircuits. The combination of anatomical accessibility, evolutionary conservation of PFC organization, and the capacity to perform complex working memory tasks with distractors establishes marmosets as a valuable complementary model to macaques for investigating the neural basis of cognitive control.

## Acknowledgements

The authors wish to thank Cheryl Vander Tuin, Whitney Froese, and Hannah Pettypiece for animal preparation and care, as well as Peter Zeman for technical assistance. We would also like to thank David Everling for assistance with touchscreen testing. This research was supported by the Canadian Institutes of Health Research (CIHR) to SE and the Canada First Research Excellence Fund to BrainsCAN. RW was also supported by the Canada First Research Excellence Fund to BrainsCAN and the Next Generation Networks for Neuroscience (NeuroNex). JS was supported by a Natural Sciences and Engineering Research Council (NSERC) Canadian Graduate Scholarship (Doctoral).

## Notes

### Competing Interest Statement

The authors have declared no competing interest.

